# Simultaneous Monitoring of Behavior & Peripheral Clocks in *Drosophila* Reveals Unstructured Sleep in an Alzheimer’s Model

**DOI:** 10.1101/013730

**Authors:** Eleonora Khabirova, Ko-Fan Chen, John S. O’Neill, Damian C. Crowther

## Abstract

Sleep and circadian rhythms are ancient, related phenomena controlled by distinct circuitry, whose appropriate regulation is critical to health. The regulatory machinery underlying sleep homeostasis is ill-defined, but biological clock mechanisms are better understood: from ‘clock gene’ oscillations to rest/activity cycles. Age- and neurodegeneration-related deterioration in sleep/wake timing was first described in humans decades ago, and has recently been recapitulated in model organisms. To delineate causal relationships between aging, sleep, neuronal function and the molecular clockwork, we developed FLYGLOW, a bioluminescence-based system which allows rest/activity, sleep consolidation and molecular rhythms to be quantified simultaneously in many individual flies over days/weeks. FLYGLOW outperforms existing methods, and allows multiparameter correlational analyses within and between flies. We also show unambiguously that peripheral tissue rhythms free-run independently of the central pacemakers driving behavioural cycles. Using an Alzheimer’s fly model we observe a profound disorganization of sleep/wake cycles, phenocopying the human disease.

## Introduction

In animals, sleep and wakefulness are regulated by both homeostatic and circadian processes (Cirelli and Bushey, 2008; Wulff et al., 2010). Sleep homeostasis is highly conserved in metazoans, but its fundamental nature is poorly understood compared with the circadian clock. This latter (about daily) biological rhythm, when synchronized with the external environment, allows an organism to anticipate, or resonate with, the varied demands of the day/night cycle, allowing metabolic and behavioral adaptions that are evolutionarily beneficial. The conservation of circadian rhythmicity across the phylogenetic spectrum from cyanobacteria to mammals indicates that these benefits are substantial (Hardin and Panda, 2013; Reddy and O’Neill, 2010). In many organisms however, these benefits are likely lost as circadian regulation of behavior and physiology progressively deteriorates with advancing age (e.g. Luo et al., 2012; Nakamura et al., 2011; Paudel et al., 2010; Rakshit et al., 2012; Tranah et al., 2011; Umezaki et al., 2012); indeed many age-related neurodegenerative disorders in humans, including Alzheimer’s Disease (AD) (Hatfield et al., 2004; Volicer et al., 2001; Wulff et al., 2010), are associated with sleep-wake abnormalities from their earliest clinical stages. Moreover a growing number of observations suggest that circadian and sleep deficits may contribute to the pathogenesis of major neurodegenerative disorders (Chen et al., 2014; Coogan et al., 2013; Long et al., 2014; Pallier et al., 2007; Park et al., 2012). Unraveling this putative reciprocal interaction between aging, neuronal (dys)function and circadian sleep/wake cycles is a non-trivial problem however, due to the challenges inherent in performing longitudinal experiments at multiple levels of biological scale in freely behaving individuals.

At a molecular level the circadian clock machinery is highly conserved from flies, through mice to humans, consisting of a series of interlocked transcription-translation feedback loops observable in every cell (Hardin and Panda, 2013). In mammals at least these rhythms are orchestrated by a central master clock mechanism located within the CNS (Mohawk et al., 2012). The resulting temperature-compensated gene expression rhythms, with an endogenous period of approximately 24 hours, are entrained to the environment *via* a number of cell-autonomous mechanisms and systemic cues, in response to exogenous signals such as light, feeding or temperature as the clock signal (*zeitgeber*). Such molecular clock oscillations in *Drosophila* may be visualized non-invasively by heterologous expression of firefly luciferase fused with a cycling ‘clock protein’, such as Period. When the organism is fed the luciferase substrate (luciferin) the level of the fusion protein is reported by the intensity of the emitted light and this signal faithfully reports the cycling of the molecular clock in every cell where it is expressed (Brandes et al., 1996; Plautz et al., 1997b).

There is much interest in mechanistic relationships within the hierarchy of intracellular, intercellular and behavioral rhythms, for example to what extent peripheral cellular rhythms are merely slave oscillators, tightly coupled to a master central clock which also co-ordinates activity rhythms (Giebultowicz et al., 2000; Hege et al., 1997; Krupp et al., 2013; Mohawk et al., 2012; Myers et al., 2003; Saini et al., 2013; Xu et al., 2008). Our ability to study the relationships between these various oscillations has until now been limited by the need, at least in *Drosophila*, to study molecular and behavioral rhythms in separate groups of organisms. As a result our observations of circadian oscillations are population-based, measuring behavioural or molecular markers not both, and report group parameters such as period, amplitude and rhythm quality with inter-individual differences being reported largely in terms of variance.

Behaviors such as locomotor activity and sleep are readily observable in a wide range of animals. In *Drosophila*, sleep is distinguished from wakefulness by a decreased sensitivity to arousing stimuli and homeostatically regulated rebound following sleep deprivation (Gilestro et al., 2009; Shaw et al., 2000). For practical purposes fly sleep is defined as a period of immobility exceeding five minutes (Gilestro, 2012; Gilestro et al., 2009).

In this study we describe for the first time a technique that allows analysis of the relationships between peripheral tissue rhythms, rest/activity cycles (a proxy for the central clock) and sleep consolidation. The characteristics of circadian rhythms in control organisms are compared to those in flies expressing the toxic Aβ peptide as a model of AD. We do this by making simultaneous real-time measurements of molecular clock dynamics and multiple behaviors simultaneously, over arrays of dozens of individual flies.

## Materials and Methods

### Fly strains and husbandry

All *Drosophila* strains in this study were housed and aged on standard cornmeal food. Flies expressing the E22G (Arctic) variant of amyloid beta peptide 1-42 (Aβ_42_) are used as a model of amyloid toxicity and are described elsewhere (Chen et al., 2014; Crowther et al., 2005). To monitor clock gene expression in control or pan-neuronally Aβ_42_ expressing flies, a new fly strain (*elav-gal4;; XLG-luc2/TM3*) containing *elav-gal4*^*c*155^ driver and the *period* promoter driven Period-luciferase fusion construct, *XLG-luc2* (Figure 1A and Veleri et al., 2003), were generated and crossed to the *UAS-Aβ*_*42*_ or a background control strain. The following offspring were studied: *elav-gal4;uas-Aβ_42_/*+; *XLG-luc2/*+ and *elav-gal4;; XLG-luc2/*+. The control and UAS-Aβ_42_ flies share the same *w*^*1118*^ background and both contain *attB* sites derived from phiC31 mediated transformation (Chen et al., 2014)

**Figure 1.**
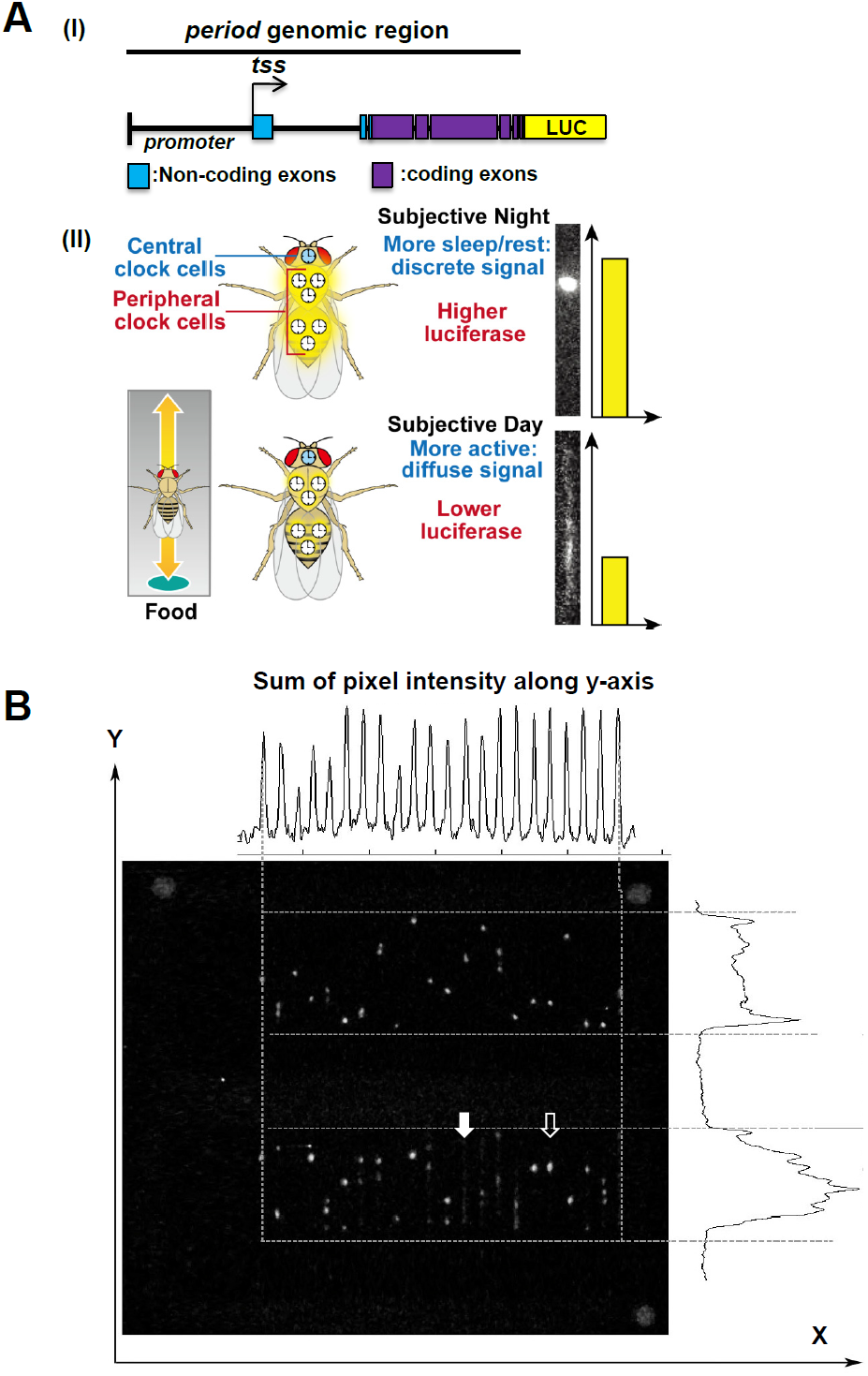
A *Period-luciferase-expressing* fly allows simultaneous measurement of the molecular clock and behavioral phenotypes. (**A**) Panel I: Period-luciferase *XLG-luc2* construct schematic containing period promoter, 5’UTR and full-length period exons and introns. Tss: transcription start site. Panel II: Stylized cartoon to illustrate that wild type flies generate a higher *Period-luciferase* signal and less locomotor activity during the subjective night, resulting in bright stationary spots in the digital image (upper section). During the subjective day the overall magnitude of bioluminescence is reduced and more evenly distributed along the length of the tube due to movement of the fly (lower section). (**B**) A representative frame from the digital camera. Three circles (grey circles) at the corners of the tray were attached as landmarks for image orientation. Total pixel intensity projections along the y-axis (upper plot) and the x-axis (right plot) were used to calculate the boundaries of each tube in pixels for all images. Active flies (filled arrow) and those at rest (blank arrow) are apparent (see also Figure 2).

### Fly arena

Individual luciferase expressing flies of required genotypes and ages were housed in a capped glass tube (cap: CAP5-Black; capillary: 5 mm × 65 mm, PGT5×65, Trikinetics. Inc. USA) containing 100 μl of 1% w/v agar, 5% w/v sucrose and 15 mM (run 1) or 50 mM (run 2) luciferin. These tubes are placed in a microscope slide storage tray (100 slide model, 165 mm × 210 mm × 35 mm, Fisher) with card dividers between the tubes (8 mm × 72 mm, 480 GSM, Ryman Limited, UK)(Figure 1-figure supplement 1A). Each tray holds up to 48 tubes. A customized arena with the mentioned dimension was constructed using black nylon plastics with fixed spacers (Figure 1-figure supplement 1B, distributed by Polygonal Tree, London, http://polygonaltree.co.uk/).

**Figure 1-figure supplement 1.**
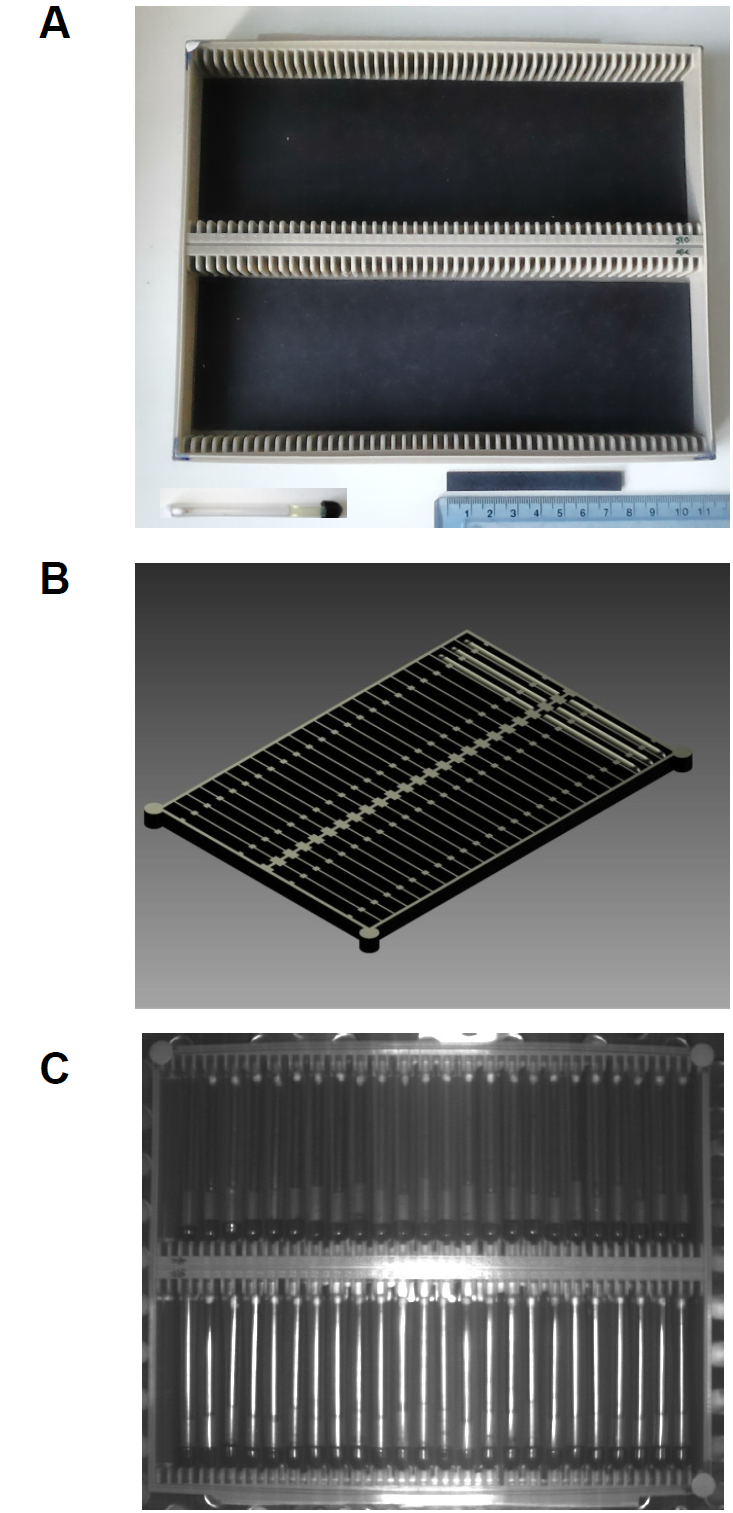
Design of fly housing arena. (**A**) Example image of plastic microscope slide tray, housing glass capillary tubes and the paper spacer. 30 cm ruler is used as scale indicator. (**B**) Illustration of the customized arena with the fixed spacer and circle markers (see Materials and Methods). (**C**) Bright field image taken by EM-CCD camera. Three paper circles (grey circles) at the corners of the tray were attached as landmarks for image rotation.

### Bioluminescence recording

Flies, 20 days post eclosion, were placed in capillary tubes and exposed to a 12-hr light: 12-hr dark (LD) regimen for three days of circadian entrainment prior to being transferred into recording conditions at anticipated dusk (ZT12). Recordings were performed under constant darkness at 26°C over seven days. Bioluminescence was detected using an EM-CCD camera (Hamamatsu Photonics UK Ltd, C9100-14) cooled to -70°C. incorporated within a Cairn Alligator system (Cairn Research Limited, UK). A bright field image was taken before each recording to ensure appropriate focus and tray alignment (Figure 1-figure supplement 1C). Bioluminescence images were recorded with contiguous 5 min integrations over 7 days with camera settings: 4× gain, 200× EM gain (Figure 1B). An example bioluminescence time series is included as a compressed movie, each frame resulting from a 5 minutes exposure (supplementary video). Week-long recordings were made with trays containing 36 control and 32 Aβ_42_-expressing flies. In separate experiments, control flies were loaded individually into the wells of a microtiter plate containing the food-luciferin substrate where their movement was restricted by covering, pierced plastic domes (Stanewsky et al., 1997). The plate was placed in the Cairn Alligator system and recorded in parallel with the tube-based assay condition. The behavior of equivalent flies was also recorded using the DAM system beam-breaking apparatus (Rosato and Kyriacou, 2006) (TriKinetics Inc., MA USA).

### Image rotation, background subtraction and feature enhancement

The raw data from the camera consisted of the 16 bit photon counts summed over 5 min at a 1024 × 1024 pixel resolution. Pre-processing of the raw data began with an estimation of the intensity of the background signal. This was achieved by averaging the brightness of pixels in a 64 × 64 square at the bottom right of the screen; this average value was then subtracted from all pixel intensities. The next step was the rotation of each frame so that the fly tubes become properly aligned with their long axes parallel to the y-axis of the image. This was achieved by manual location of three marker points (grey dots in corners, Figure 1B) on the tray in which the fly tubes are placed. Once correctly aligned the corresponding tubes in successive rows lie directly above each other on the y-axis. In order to identify where, across the image, each pair of corresponding tubes was located we plotted the profile of the pixel intensities summed along the y-axis. This profile oscillates, with each peak of the summed pixel intensities indicating where the centre of each tube can be found (Figure 1B). A similar process was performed along the y-axis, summing pixel intensities across the image to detect the top and bottom of each row of tubes. Once these steps were complete we assigned a rectangle for each tube that corresponded to its boundary, assuming that each tube was 17 pixels wide. We also assigned a rectangle that corresponds to the dark background between pairs of tubes that is centered on the midpoint between adjacent tubes and was 7 pixels wide. The average pixel intensity of this rectangle was termed the “inter-tube background intensity” and was used in the tube-by-tube background subtraction process described below. At this stage the data was saved as pre-processed “raw data” and was used for all quantitative calculations. To allow sensitive feature detection we also enhanced the contrast and brightness of the images so that flies could be reliably identified despite changes in overall brightness. This process of enhancement used the contrast-stretching transformation algorithm in MATLAB that globally optimizes the dynamic range of pixel intensities. Such processed images were stored as “enhanced data”.

### Detecting resting and moving flies

Hardware issues result in small systematic variations in pixel brightness between regions of the image (vignette effect). To compensate for this we performed a second round of background intensity subtraction. This was done on a tube-by-tube basis using the inter-tube background intensity, calculated as above. We then divided each tube rectangle into 4-pixel-high bins along their long axis (y-axis). We summed pixel intensities for each bin and calculated the mean and standard deviation of these values. Bins containing a resting fly were characterized by peaks in the bin intensity profile along a tube (Figure 2, “Enhanced images”); a bin contained a resting fly when maximum pixel intensity of the peak > (Mean tube intensity + Standard Deviation).

**Figure 2.**
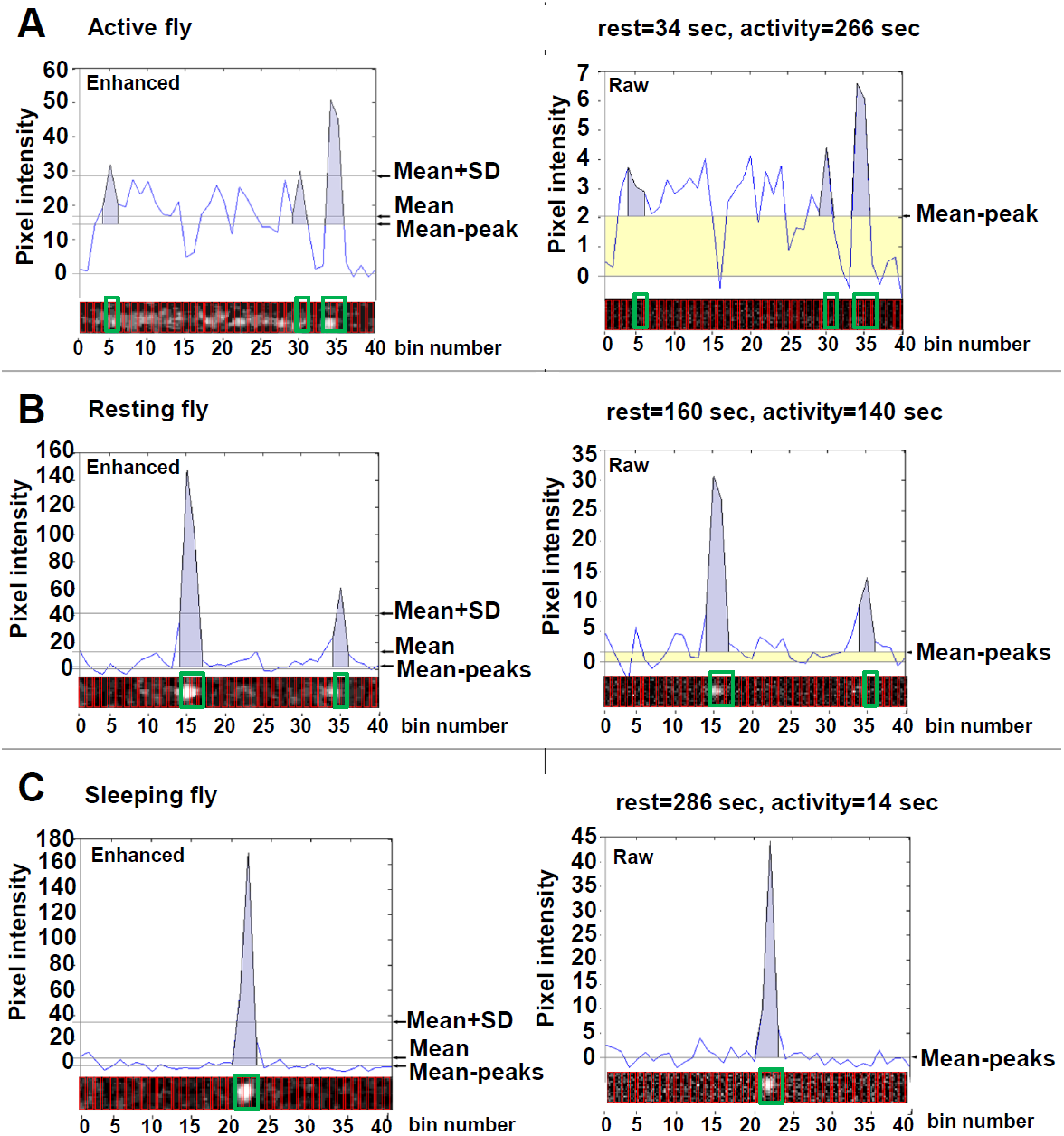
Frame-by-frame measurements of molecular clock and behavioral status. Pixel intensity was plotted and binned into 40 segments along the length of the tube for each image (red boxes). The enhanced images (left column) permitted sensitive feature detection and were used to detect bins containing peaks of intensity: The mean pixel intensity and standard deviation (SD) for all bins in a tube were calculated. Peaks in the enhanced data were assigned when the intensity of a bin exceeded the mean+SD for that tube. Using the raw images the value of the mean bin intensity, excluding bins containing peaks (green boxes), was calculated (the Mean-peak value). The yellow area below this value represents the light emitted by moving flies. The purple area under the peaks represents resting flies. The total of these areas is equivalent to 300 sec. Representative data is presented for frames containing an active fly (**A**), an active fly exhibiting periods of rest (**B**) and a sleeping fly (**C**). The total pixel intensity within a particular tube indicated the level of the *Period-luciferase* construct and hence the status of the molecular clock.

We then excluded bins that contained resting flies and used the raw data from the remaining bins to recalculate the mean tube intensity (Mean without peaks, Figure 2). For each fly and each frame we partitioned the 300 sec of the camera exposure time between two parameters: i) time spent active and ii) time spent resting. The contribution of activity to the profile was measured by calculating the area of the rectangle between the (Mean without peaks) and the background (Figure 2, yellow areas). The contribution of resting to the profile was measured by calculating the area under each identified peak, bounded by the (Mean without peaks) (Figure 2, purple areas).

For each frame:

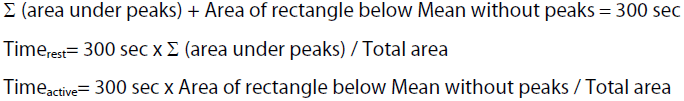

In fewer than five percent of frames, subtraction of the inter-tube background resulted in a negative value for a tube’s mean intensity; this was usually due to artifactually high intensity pixels in the background region. In these circumstances the data was discarded and the values of Time_rest_ and Time_moving_ interpolated from adjacent time points.

### Measuring the oscillation of the molecular clock

The oscillation of molecular clock was calculated by summing up the background-adjusted pixel intensity in a rectangle constrained by each tube’s coordinates. This primarily reports cellular circadian rhythms of gene expression in peripheral tissues of the fly.

### Assigning sleep episodes

*Drosophila* sleep episodes are defined as a period of locomotor inactivity lasting for >300 sec. Having calculated the time spent moving and resting for each frame we defined the presence of a sleep episode in the following way: Firstly, a sleep episode can only be initiated in a frame with a single intensity peak, indicating that the fly was resting in only one position. Secondly, considering the subsequent frames in turn we checked that there was a peak at the same position, indicating that the fly had rested from one frame to the next. Subsequent frames were analyzed sequentially until the resting peak was lost. The component rest times were then summed across the multiple frames; when the total resting time exceeded 300 sec then the episode was defined as sleep. This approach is conservative and does not over-identify brief rests as sleep episodes. Each frame within a sleep episode was assigned a 1 in the binary sleep array; all other frames are assigned a 0 (1=asleep and 0=awake). Using this method, we can robustly assign one of three behavioral states to each frame for each fly; these are “active” (Figure 2A), “active with rest” (Figure 2B) and “sleeping” (Figure 2C).

### Calculating sleep consolidation

Our novel sleep consolidation index provides an indication of how long a block of sleep lasts. To calculate this index for a particular fly we looped through the binary sleep array and identified blocks of sleep as consecutive episodes separated by wake periods. Each time point within a particular sleep block was assigned with the duration of its encompassing sleep episode (Figure 3).

**Figure 3.**
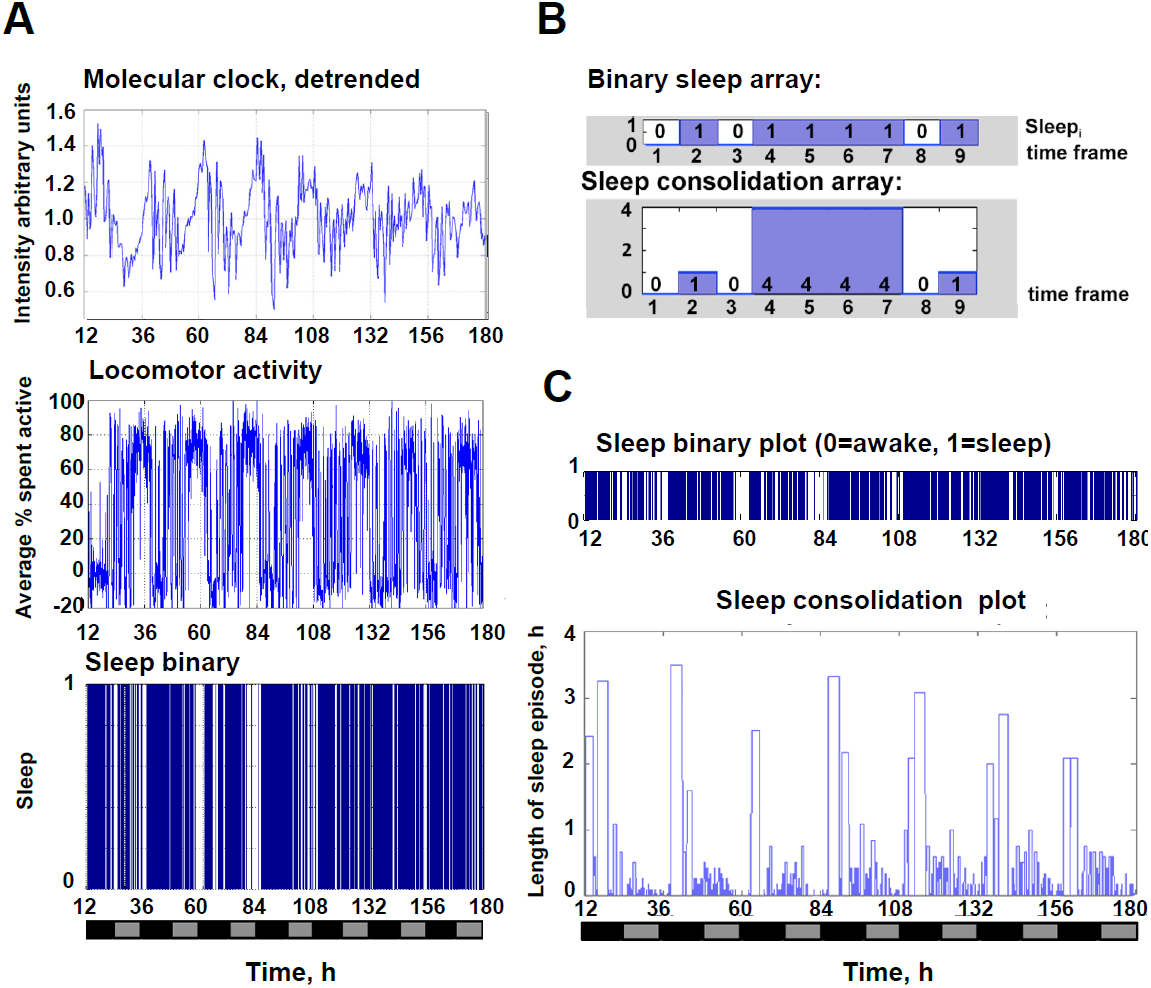
Time course of molecular and behavioral measurements for a single fly. (**A**) The bioluminescence intensity of a representative single fly during seven days of constant darkness reported the oscillation of the molecular clock (Molecular clock). The percentage of time spent active (Locomotor activity) was calculated and the sleep status (Sleep binary) was assigned to a frame as described in the Materials. (**B**) Each frame containing a fly that had been immobile for over 300 sec was assigned a value of 1 in the binary sleep array. All other frames were assigned a value of 0. The sleep consolidation array contained, for each sleep frame, the total length of the encompassing sleep episode. (**C**) This measurement of sleep consolidation exhibited circadian oscillation. X-axis in (A) and (C) indicates circadian time in hours (Time, h) with subjective day (grey) and night (black) indicated. The recording started from subjective dusk (circadian time=1200 hr).

### Final data output

The temporal resolution of the data was reduced to 30 min by binning data from consecutive timepoints, similar to the conventional circadian locomotor assay (Chen et al., 2014). For time spent resting or moving, the binned value represented the sum of the constituent timepoints. For sleep consolidation, the binned value was assigned as the mean value of non-zero entries in the sleep consolidation array.

### Time-series analysis

We adapted the well-established autocorrelation methodology to determine rhythmicity and circadian period using Flytoolbox in the MATLab environment. To make our analysis comparable to the DAM actimetry system we have limited our observations to 7-days with data considered in 30 min bins. This constrained paradigm permits the convenient use of Rhythmicity Statistic analysis as described by Levine at al. (Levine et al., 2002a). To normalize the exponential decay of bioluminescence intensity with time we employed the detrend filter. We also combined the sliding-window method (3 day window) with the phase determination tool (peakcircleplot, Levine et al., 2002a; Levine et al., 2002b) to quantify differences in circadian phase between time series.

## Results

### Simultaneous molecular clock and behavioral measurements: FLYGLOW

Our experimental *XLG-luc2* flies (Figure 1A, I) express a Period-Luciferase fusion protein and produce bioluminescence when fed with luciferin (Figure 1A, II). This light signal is detectable using a sensitive electron-multiplying CCD camera, and effectively reports clock gene expression in peripheral tissues, that is lowest at anticipated dawn (CT0) and highest at anticipated dusk (CT12) (Figure 1A). When constrained within glass capillary tubes (Figure 1A) these flies are considered to exhibit one-dimensional locomotor behavior that is readily quantified by computational image processing. In these experiments one end of the tube was sealed with fly food containing luciferin and the other end closed with cotton wool; these conditions will support a *Drosophila* for more than three weeks. By integrating the bioluminescence across 300 sec exposures for each frame (Figure 1B) we constructed a time-lapse movie representing 7 days of fly behavior (supplementary movie). For the area represented by each tube, within each frame there is sufficient information to directly measure the level of peripheral clock gene expression reported by luciferase activity (number of photons detected *per* region of interest) as well as locomotor activity (spatial distribution of detected photons within that region); from the latter we can reliably infer the duration of fly sleep episodes according to the widely accepted standard (Gilestro, 2012; Gilestro et al., 2009).

In brief the images were processed to detect two patterns of intensity along the length of each capillary tube. Active flies appeared as a smear along the tube (Figure 1A, B, filled arrow) and generated a rectangular area on the intensity profiles (Figure 2, yellow shading). Resting flies generated local bright spots (Figure 1A, B, empty arrow) that were detected as peaks on the profile (Figure 2, purple shading). Contrast-enhanced images were used to detect the bright spots of resting flies; thereafter, raw data, processed only to subtract local background camera signal, was used for all quantitative measures. For each 300 sec frame and for each fly our analysis partitioned time according to the area under the “mean without peaks” value on the intensity profile (yellow) versus the area under the peaks (purple). The sum of all the yellow and purple segments was equivalent to 300 sec for each frame and for each fly.

Because the absolute intensity of the bioluminescence decreased exponentially with time, due to decay of the luciferin, the data was detrended so that the signal had the same mean and amplitude across the 7-day time course (Levine et al., 2002a, b). This peripheral molecular clock signal was then plotted for the population alongside contemporaneous observations of frame-by-frame quantification of activity and sleep (Figure 3A). The conventional sleep binary plot (Figure 3A, sleep binary) depicted total sleep along the time, but failed to demonstrate the sleep episode fluctuation. Further processing of the data allowed a measure of sleep consolidation to be generated (Figure 3B) and facilitated the visualization of circadian oscillation in sleep episode length. The sleep consolidation plot indicates that sleep episodes of over 3 hrs are commonplace in the early part of the subjective night (marked with black bars on x-axis, Figure 3C). Later in the night discrete episodes of 40-50 min are seen but by contrast during the day only shorter naps are seen (<30 min). This pattern is the first clear visualization that *Drosophila* shows daily fluctuation in sleep quality and is reminiscent of the structure of human sleep where deep, slow wave sleep is apparent particularly in the early part of the night whereas rapid eye movement (REM) sleep predominates towards morning (Wulff et al., 2010).

### Comparison of FLYGLOW with existing experimental paradigms

We then compared the actimetric performance of FLYGLOW with the widely-adopted DAM beam-break-counting apparatus (Rosato and Kyriacou, 2006) (TriKinetics Inc., MA USA). We used two populations of flies, the first were strongly rhythmic control flies and the second pan-neuronally expressed toxic amyloid β peptide - a model for Alzheimer’s disease (Figure 4A). Previous work has shown that the Aβ expressing flies show progressive behavioral dysrhythmias as they age despite their central molecular clock remaining relatively intact (Chen et al., 2014). Considering the data for control flies (left panels), as expected the DAM system underestimates locomotor activity as compared to FLYGLOW. In the DAM profile the underlying oscillation in behavior has relatively low amplitude with narrow bursts of beam breaking. By contrast the FLYGLOW profile is less noisy with higher amplitude rhythmic signal, since it misses far fewer false negatives i.e. movement without beam-breaking (consistent with a recent video tracking system, Gilestro, 2012). The Aβ-expressing flies exhibit markedly weaker rhythmicity as compared to controls (right panels), appearing essentially arrhythmic in the DAM system but retaining detectable oscillation in FLYGLOW. The quality of each fly’s behavioral rhythmicity may be quantified by calculating a time-delayed auto-correlation statistic (RS value, Levine et al., 2002a). In this case we saw that both DAM and FLYGLOW rank Aβ-expressing flies as less rhythmic than controls (p<0.001 for both techniques, Figure 4B). Notably however the RS values are higher overall for FLYGLOW as compared to DAM (p<0.05) indicating that the new method may be more sensitive at detecting behavioral rhythms than the existing approach. When we examined the distribution of RS values in the population of flies we see that the ability of FLYGLOW to detect very highly rhythmic flies accounts for the bulk of the performance gain (Figure 4C).

**Figure 4.**
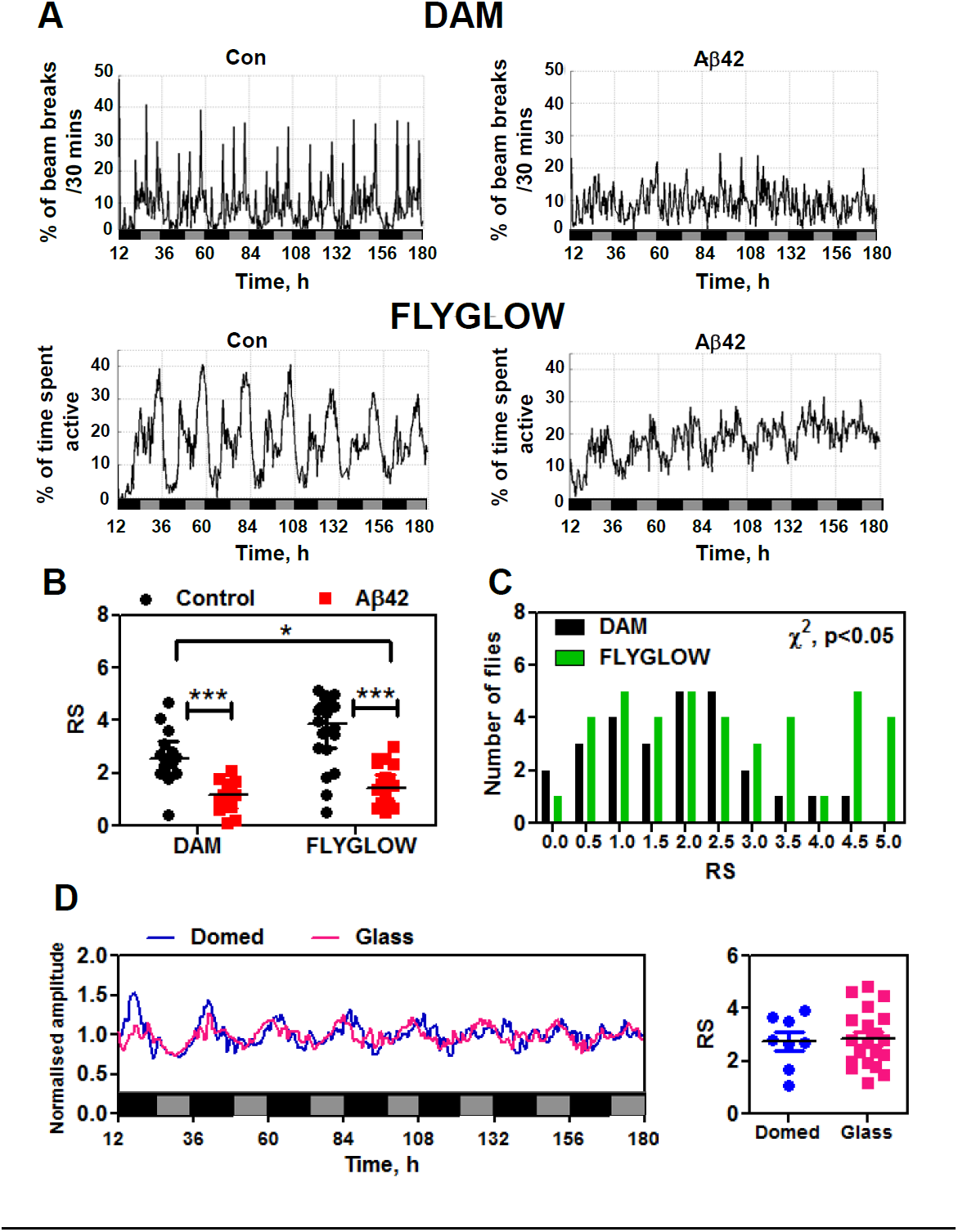
Comparison of FLYGLOW with conventional methods for population measurements of locomotor activity and molecular clock rhythms. (**A**) The mean locomotor activity for control (n=14) and Ap-expressing flies (n=13) was assessed using the DAM system (y-axis: percentage of the maximum beam breaks per 30 minutes) and FLYGLOW (n=20 for control and n=22 for Ap-expressing flies). The x-axis represents circadian time in hours (Time, h) with subjective day (grey) and night (black) indicated. The recording started from subjective dusk (circadian time=1200 hr). (**B**) The robustness of locomotor rhythmicity was determined by calculating RS values. Rhythms were significantly stronger in the control flies as compared to Aβ-expressing flies (2-way ANOVA, p<0.001). The RS value of the FLYGLOW data as a whole was higher than that of DAM (2way ANOVA, p<0.05). (**C**) The frequency distribution of RS values from FLYGLOW (n=42) is significantly different compared with those from DAM (n=27) (χ^2^-test, p<0.05). FLYGLOW was particularly sensitive at detecting highly rhythmic locomotor behavior (RS>4.0). (**D**) Left panel: Comparing the mean bioluminescence rhythms from FLYGLOW (n=20, magenta) with flies observed in domed 96-well plates (n=8, blue) indicates that the molecular clock signals are essentially identical. Right panel: There are no significant differences between RS values of the molecular clock rhythms between FLYGLOW (magenta) and the classical approach (blue).

Next we compared FLYGLOW molecular clock detection with the performance of the existing approach in which flies are constrained under small plastic domes within the wells of a 96 well microtiter plate (Stanewsky et al., 1997). We compared our new approach with this current method to exclude the possibility that the extra freedom of movement seen in FLYGLOW could add a systematic artifact of the system: for example the availability of the luciferin substrate might be less in the tubes and result in artifactual changes in bioluminescence. As shown in Figure 4D, the two approaches generate essentially identical bioluminescence traces; moreover the fly-by-fly rhythmicity analysis indicates that the RS values are also very similar.

### FLYGLOW simultaneous comparison of molecular and behavioral oscillations

Three principal datasets are routinely generated by the FLYGLOW system; these are, i) the molecular clocks in peripheral tissues as reported by total bioluminescence, ii) percent of time spent in locomotor activity and iii) sleep consolidation. These are conveniently presented as mean data for a population of flies (Figure 5A). Control flies aged 23-30 days (left column, n=36) show high amplitude molecular oscillations with gradual dampening (Veleri et al., 2003) that peak in the early part of the subjective night and retain their circadian rhythm throughout the 7-day experiment. For flies expressing the Aβ peptide (right column, n=32 flies) the molecular rhythms are also well preserved although there is more deterioration from day 4 onwards.

**Figure 5.**
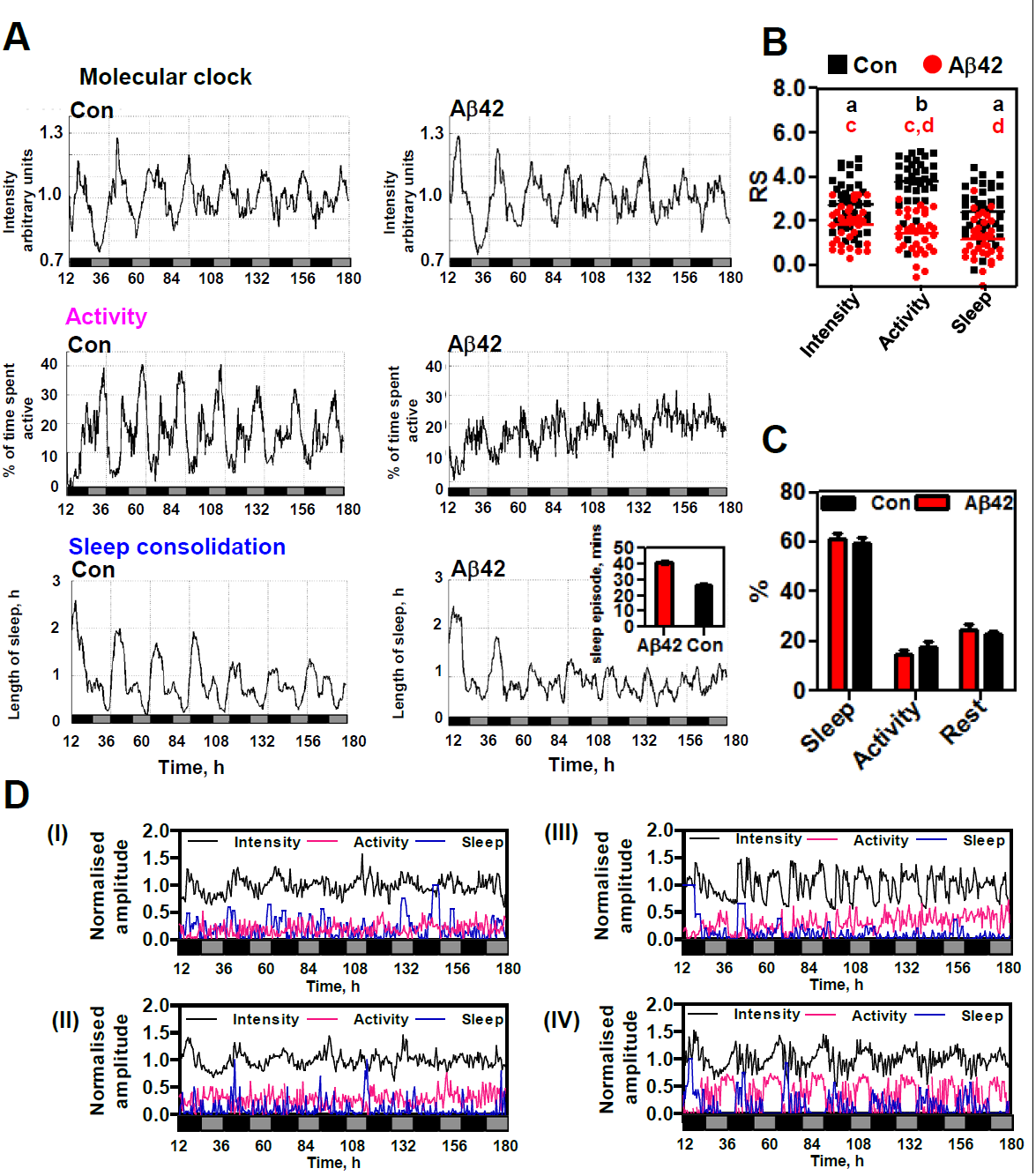
Comparing the rhythmicity of control and Aβ-expressing flies. (**A**) The population means of *XLG-luc2* signal intensity (Molecular clock), percentage of time spent active (Activity) and sleep episode length (Sleep consolidation) were plotted for control (n=36), and Aβ-expressing (n=32), flies. The insert indicates that daytime sleep episodes were longer in Aβ-expressing flies (red) as compared to controls (black). (**B**) All three rhythms are significantly less robust in Aβ-expressing flies (red dots) as compared to controls (black squares) as determined by comparison of RS values by matched two-way ANOVA. Repeated measures one-way ANOVA was used to identify statistical differences in RS values for various pairs of rhythms within each genotype. The same alphabets are indicated for the group with no difference in the RS values (p>0.05), e.g. the average RS value of the intensity rhythms (a) in the control flies are the same to those of the sleep rhythms (a), but are different from those of the activity rhythm (b). (**C**) Despite the loss of rhythmicity seen for Aβ-expressing flies, there were no significant differences in the relative partitioning of time between sleep, activity and rest (p>0.05, by χ^2^-test or matched 2-way ANOVA). (**D**) Panels I & II: Individual recordings for Aβ-expressing flies with weak behavioral rhythms (activity in magenta and sleep consolidation in blue) and more robustly rhythmic molecular clock oscillations (black). Panel III: representative Aβexpressing fly with non-rhythmic molecular clock and behavioral signals. Panel IV: representative control fly with robust molecular and behavioral rhythms. Sleep consolidation amplitudes are normalized to the longest sleep episode. X-axis in (A) and (D) indicates circadian time in hours (Time, h) with subjective day (grey) and night (black) indicated. The recording started from subjective dusk (circadian time=1200 hr)

For control flies, the percent of time spent in activity is also highly circadian. In these controls FLYGLOW can clearly distinguish two activity peaks, the first occurring just before subjective dusk, when flies spend up to 40% of their time moving, and the second just before dawn when they are moving approximately 30% of the time. The locomotor behavior for Ap-expressing flies exhibits a low-amplitude, less regular rhythm that can be characterized as a state of continuous restlessness throughout the subjective day and night.

As expected, the sleep consolidation plot is highest when the locomotor activity is least, with at least a superficial resemblance to the deep sleep (non-REM) that humans tend to manifest earlier in the night. Specifically, the duration of the average sleep episode in the early night, when control flies spend <5% of their time moving, is remarkably high, at more than 2 hrs. During the morning of the subjective day the flies take short naps of up to 30 min at a time when they exhibit significant locomotor activity (20-30% time spent moving). Sleep episodes are at their shortest during the late afternoon peak in locomotor activity. The inset indicates that Aβ-expressing flies (red), similar to patients with AD, have longer sleep episodes than control flies (black) during the subjective day.

Our qualitative interpretation of population behaviors (Figure 5A) was confirmed objectively by calculating rhythmicity statistic values (RS values). We found that for all three datasets the RS values are significantly higher in control flies than in their Aβexpressing counterparts (Figure 5B). Remarkably these changes in rhythmicity do not cause any statistically significant difference in the relative proportions of sleep (sum of rests longer than 5 min), activity and rest (sum of rests shorter than 5 min) (Figure 5C). Thus the differences in sleep relate entirely to its temporal organization and not its total amount.

The plotting of representative data from individual flies may also be instructive (Figure 5D). For example, many Aβ flies exhibit chaotic behavior (panels I & II) however it is not uncommon in this context to see ongoing molecular rhythms. At one extreme an individual Aβ fly may loose rhythmicity in all three signals (panel III) while the majority of control flies remain rhythmic by all three parameters throughout a week long recording (panel IV). Using paired statistics, we found that behavioural rhythms showed greater deterioration in individual Aβ flies than molecular oscillations (Figure 5B).

### Dissecting the mechanistic links between the various rhythms

Our single fly measurements permit powerful analysis of correlations between the quality and the period of the three rhythmic datasets. Indeed the simultaneous observation of multiple rhythms within a single fly allows the use of paired statistical tests, something that is not possible when comparing separate population-based measurements. Considering rhythm quality, if the RS values of two rhythms are correlated then one may conclude they are coupled, that is that they share a common clock mechanism. For example when one of the rhythms is robust in a particular fly one expects that the other will also be similarly robust, and vice-versa. Our systematic three-way analysis of RS correlations in control flies showed that there is no evidence for coupling between the peripheral molecular clock and either of the two behavioral phenotypes (Figure 6A: activity, panel I & sleep, panel II). By contrast the RS values of the two behaviors were correlated (panel III, p<0.05), indicating that locomotor and sleep rhythms are both influenced by the same central clock mechanism, but presumably *via* different output pathways.

**Figure 6.**
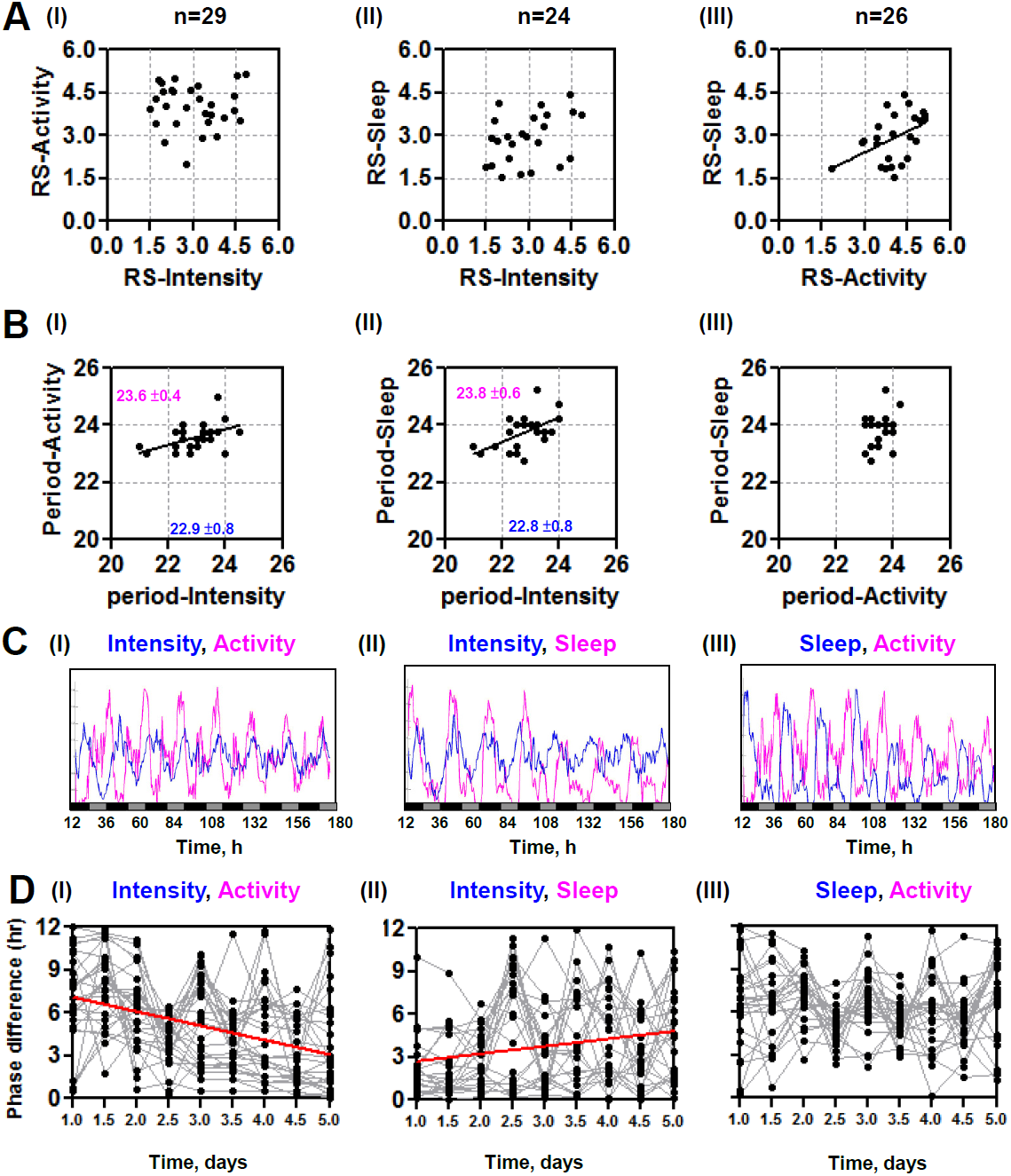
RS and period correlation analysis: dissecting causal relationships. (**A**) The paired RS values for the various rhythms were plotted for each fly. Using only control flies there was no correlation between RS values for molecular clock (Intensity) and locomotor activity (panel I) or sleep consolidation (panel II). By contrast the RS values for locomotor activity (Activity) and sleep consolidation (Sleep) rhythms where correlated (panel III, p<0.05). Only flies rhythmic for both tested signals were considered, resulting in varying n-values between panels. (**B**) The paired signal periods for the various rhythms were plotted for each fly. Using only control flies we found that the periods of locomotor activity (panel I) and sleep consolidation (panel II) rhythms were both correlated with the period of the molecular clock (both p<0.05). By contrast there was no correlation between the periods of the sleep consolidation and locomotor activity rhythms (panel III). The average periods of the behavioral rhythms (magenta) both differed significantly from the shorter molecular clock period (blue). (**C**) The population data for the corresponding pairwise rhythm comparisons were plotted. The molecular rhythm had a shorter period than either behavior rhythm (panels I & II) however the respective behavioral rhythms had very similar periods and retained a fixed phase relationship (panel III). X-axis indicates circadian time in hours (Time, h) with subjective day (grey) and night (black) indicated. The recording started from subjective dusk (circadian time=1200 hr). (**D**) Pairwise observations also allowed the calculation of phase differences between rhythms across the time course. There is a systematic reduction in phase difference as the molecular rhythm “catches up” with locomotor activity (panel I); by contrast the phase difference increases as the molecular rhythm marches away from the sleep consolidation rhythm (panel II). There is no overall change in phase difference when comparing the two behavioral signals (panel III). The phase of each rhythm was calculated for a 3-day window. The start times of each window are separated by half a day. The grey lines connect the half-daily estimates of phase difference for each fly. The red lines indicate linear regressions that have a gradient that is significantly different to zero (p<0.01). The significant of the RS correlations was determined by the *Pearson r* with p<0.05 (A, B). The significance of the correlation between period differences *Paired t-test* with p<0.01 (B).

A similar three-way comparison of rhythm periods in individual flies, revealed correlations between the peripheral molecular clock and both of the behavioral rhythms (Figure 6B, panels I & II). This is despite the fact that the RS values are not correlated and the periods of the rhythms are not equal. That their periods *are* correlated implies that these oscillations share underlying mechanisms because, in the context of slight variability between individuals, factors that modify the period of one rhythm tend to also vary the period of the correlated rhythm i.e. they are co-variant. Strikingly, the periods of the sleep and activity rhythms do not correlate (panel III), indicating that sleep rhythms are influenced by additional mechanism such as homeostatic regulation and possibly even local brain oscillations in the fly arousal/sleep centers that are not tightly coupled with the central clock neurons (e.g. Kunst et al., 2014; Wu et al., 2008).

### The peripheral molecular clock and behaviors progressively desynchronize in the dark

While the periods of the behavioral rhythms and peripheral bioluminescence rhythm are correlated, surprisingly they are not equal (Figure 6B I & II; average periods for behaviors in magenta, 23.6-23.8 hr & molecular clock in blue, 22.8-22.9 hr). Because the molecular clock oscillates faster than either of the behaviors in constant darkness, we observed the marching of this rhythm across the behaviors during the course of the experiment (Figure 6C I & II). In these circumstances, when we plotted the phase difference between the molecular clock and activity (Figure 6D, panel I) we saw a progressive reduction in phase difference as the molecular signal caught up with the behavior. Likewise (panel II) the phase difference increased as the molecular clock progressively moved ahead of the sleep rhythm. By contrast the sleep and activity rhythms had the same period and so maintained a stable phase relationship (panel III).

## Discussion

Not only do circadian rhythms have profound roles to play in biology, they are also recognized as important factors in human disease. On one hand environmental disruption of circadian rhythms increases the risk of, for example, obesity and diabetes while on the other hand some diseases are accompanied by circadian deficits from an early stage (Rakshit et al., 2014; Reddy and O’Neill, 2010). Neurodegenerative disorders, not least AD, are characterized by sleep-wake abnormalities, indeed it is night-time confusion and wandering that often makes institutional care inevitable in such cases (Wulff et al., 2010). Considering a recent finding that any use of (benzodiazepine) sleeping tablets is linked to a 50% increase in AD risk, it is apparent that circadian deficits become symptomatic early, as part of a dementia prodrome (de Gage et al., 2014). As AD progresses, patients will exhibit marked sleep fragmentation and loss of the normal day/night partitioning of sleep. A consequence of this can be a state of continual restlessness, often worse in the early evening when the syndrome is termed sundowning (Coogan et al., 2013; Hatfield et al., 2004; Volicer et al., 2001).

*Drosophila melanogaster* is a model organism that phenocopies key aspects of the deterioration in circadian/sleep organization associated with human aging and neurodegeneration. Our inability to longitudinally monitor multiple behavioural and molecular markers in individual animals however, has prevented the full potential of the fly as screening tool from being realized as researchers try to understand the etiology of complex diseases such as AD.

We have therefore developed FLYGLOW in order to achieve our goal of understanding the causal relationship between molecular clock oscillations, behavioral and sleep rhythms, in health and disease. Our advance relies upon the synergistic optimization of multiple existing technologies. Specifically we used an exquisitely sensitive EM-CCD camera mounted upon a light-tight, environmentally controlled cabinet. The camera collected exposures of a duration that matches the current working definition of fly sleep, and thus by looking at the distribution of the light signal, per tube per exposure, fly movement during each 5 minute window could be robustly inferred. FLYGLOW also relies upon a fly strain expressing a particularly bright, peripherally-expressed clock gene::luciferase reporter, and fed very high luciferin concentrations which are necessary to sustain bioluminescence during sleep (when flies do not eat). Finally the development of new mathematical tools for the analysis of this type of time lapse imaging was essential in allowing us to explore the concept of sleep consolidation in a fly model for the first time. Our spatial tracking combined with measurement of total bioluminescence facilitated the quantification of multiple phenotypic parameters from an array of individual organisms allowing us to assign simultaneous molecular, locomotor and sleep events over a 7 day period. Our approach compares very favorably with current alternative technologies, that are limited by only measuring one of these parameters in any given experiment (sleep, activity, or bioluminescence). This has forced researchers to compare populations of flies, rather than individuals and thus loses the power of intra-individual correlational analyses over time. Importantly FLYGLOW could equally be employed for more broadly based behavioral genetic and drug-based screens, beyond sleep/wake or neurodegeneration research.

Our initial observation was that, as a population, the flies show complex patterns of sleep and locomotor behavior that remain synchronized in the constant dark. Sleep consolidation reached remarkably high levels in the early night, with episodes of 2-3 hours being commonplace. In flies expressing the Aβ_42_ peptide this high quality “deep” sleep was lost early and instead was partitioned more uniformly across the day, resulting in longer daytime naps. These changes in sleep structure strikingly resemble the deficits seen in patients with AD (Coogan et al., 2013; Wulff et al., 2010). Interestingly there was no overall change in the relative amounts of rest, sleep and activity in the control and Aβ_42_-expressing flies indicating a change in the structure rather than the amount of sleep between the two groups.

Remarkably, the molecular clock in peripheral tissues was more robust than its behavioral counterparts. Echoing our previous observations of the central molecular clock becoming uncoupled from behavior in Aβ_42_-expressing flies (Chen et al., 2014), this study also revealed many examples of flies exhibiting robust molecular oscillations contemporaneously with behavioral arrhythmia. This recurring observation of molecular-behavioral dissociation suggests that one approach to re-entraining disturbed behavioral rhythms may be to enhance the output of the central neuronal clock.

An important conclusion of this study is that single-fly observations of multiple circadian phenotypes allow us to use RS and period correlations to dissect the causal relationships between circadian oscillations. The ability to use paired statistical tests greatly increases the power of such comparisons. Despite this, we observed no correlation between the RS values for the peripheral molecular clock and either of the behavioral rhythms. This confirms that there is no global peripheral molecular-behavioral coupling in our system as suggested previously by a number of investigators (Giebultowicz et al., 2000; Hege et al., 1997; Plautz et al., 1997a). The contrasting finding that RS values for sleep consolidation and locomotor behavior are indeed correlated indicates that these rhythms are generated by a shared central clock.

Similar studies of period correlations between rhythms have allowed us to detect oscillations with shared mechanisms despite their having no direct coupling. Thus our observation that there is a period correlation between the peripheral molecular clock and both behaviors is more notable because not only do the periods differ by almost one hour but also because there is no corresponding RS correlation. A corollary of this finding, and one confirmed experimentally, is that the molecular and behavioral signals desynchronise in the dark as the shorter period molecular oscillation marches across the behavioral rhythms. The intriguing absence of a period correlation between sleep and locomotor activity points to more complex regulation of sleep/wake that may respond to other homeostatic and regulatory signals (Cirelli and Bushey, 2008; Gilestro et al., 2009; Myers et al., 2003; Naidoo, 2009). Similar dissociation occurs in humans during forced desynchrony (14 hr:14 hr, L:D) when circadian physiology becomes decoupled from sleep-wake cycles. Such observations however have never been possible in *Drosophila* before.

**In conclusion we present FLYGLOW: a new broadly applicable approach that for the first time allows the causal relationships between molecular and behavioral circadian rhythms to be dissected by simultaneous molecular and phenotypic observations in individual flies. Since the running costs of our method are low, but is amenable to high-throughput genetic screens, we expect it to furnish novel insights into the etiology of complex age-related human diseases such as Alzheimer’s disease.**

## Acknowledgements

We thank Ned Hoyle (LMB, Cambridge) and Florian Hollfelder (Biochemistry, Cambridge) for fruitful discussion and Simon Bullock (LMB, Cambridge) for his assistance. Jo Westmoreland of MRC LMB Visual Aids Department. E.K. is supported by Wellcome Trust 4-year PhD programme in Mathematical Genomics and Medicine. J.S.O’N. is supported by the Medical Research Council (MC_UP_1201/4) and the Wellcome Trust (093734/Z/10/Z). D.C.C and K.-F.C. are supported by the Wellcome Trust (082604/2/07/Z). D.C.C. was an Alzheimer’s Research UK Senior Research Fellow (ART-SRF2010-2).

## Author contributions

D.C.C., K.-F.C. and J.S.O’N. designed the FLYGLOW strategy. K.-F.C. and J.S.O’N. performed the experiments. E.K. built the image processing/analysis pipeline. K.-F.C., E.K., J.S.O’N., and D.C.C. analyzed the data. D.C.C. and K.-F.C. wrote the manuscript with crucial inputs from E.K. (Bioinformatics) and J.S.O’N. (Circadian Biology). D.C.C. conducted the project.

